# Metabolic state of human blastocysts measured by fluorescence lifetime imaging microscopy

**DOI:** 10.1101/2021.07.09.451821

**Authors:** Marta Venturas, Jaimin S. Shah, Xingbo Yang, Tim Sanchez, William Conway, Denny Sakkas, Dan J. Needleman

## Abstract

Mammalian embryos undergo large changes in metabolism over the course of preimplantation development. Embryo metabolism has long been linked to embryo viability, suggesting its potential utility in Assisted Reproductive Technologies (ART) to aid in selecting high quality embryos. However, the metabolism of human embryos remains poorly characterized due to a lack of non-invasive methods to measure their metabolic state. Here, we explore the application of metabolic imaging via fluorescence lifetime imaging microscopy (FLIM) for studying human blastocysts. We use FLIM to measure the autofluorescence of two central coenzymes, NAD(P)H and FAD+, in 215 discarded human blastocysts from 137 patients. We find that FLIM is sensitive enough to detect significant metabolic differences between blastocysts. We show that the metabolic state of human blastocysts changes continually over time, and that variations between blastocyst are partially explained by both the time since fertilization and their developmental stage, but not their morphological grade. We also observe significant metabolic heterogeneity within individual blastocysts, including between the inner cell mass and the trophectoderm, and between the portions of hatching blastocysts within and without the zona pellucida. Taken together, this work reveals novel aspects of the metabolism of human blastocysts and suggests that FLIM is a promising approach to assess embryo viability through non-invasive, quantitative measurements of their metabolism.

## Introduction

Mammalian preimplantation embryos undergo dynamic metabolic changes necessary to support distinctive developmental events ^1–3^. Early stages of preimplantation embryo development rely primarily on pyruvate for energy generation^3,4^. It was previously thought that once the embryo reached the 8-cell stage, the embryo underwent a metabolic switch and glycolysis became the predominant pathway for ATP production^3,5,6^. However, a number of studies have shown that glucose does not only contribute to energy production in mouse embryos, but instead plays crucial signaling roles required for blastocyst formation and cell fate specification^7–9^. Glucose is also partially converted to lactate, which is thought to facilitate different aspects of implantation, including tissue invasion, angiogenesis and modulation of the immune response^10^. In parallel, oxygen consumption increases at the blastocysts stage^10–12^. Proper metabolic function is essential for producing developmentally viable eggs and embryos^4,5,13^, and impaired metabolic activity has been correlated with decreased developmental potential in mouse^4,14^ and human embryos^4,13,15^. Embryos with the highest developmental potential were found to have low amino acid turnover ^16^ and low rates of oxygen consumption^11,17^. These results have led to the ‘Quiet embryo hypothesis’ by Leese et al. which proposes that embryos with a higher developmental capacity are characterized by a quieter metabolism rather than an active one^18^, which is suggested to be linked with cellular or molecular damage^2,19^. However, in apparent contradiction with this hypothesis, some studies indicate that pyruvate^20^ and glucose uptake^21^ levels are higher in embryos that develop successfully.

Blastomeres in preimplantation embryos are genetically, morphologically and metabolically heterogeneous. Metabolic heterogeneity between blastomeres is thought to be associated with either cell lineage specification^7^ or cellular stress^16^ as high levels of heterogeneity have been associated with developmental arrest^22^. Cellular stress may be associated with increased DNA damage ^3^ which could be either a cause or a consequence of cell metabolic (dys)function. At the blastocyst stage, clear cellular differentiation already exists, with the apical trophectoderm (TE) and inner cell mass (ICM), which give rise to the placenta and fetus, respectively. Studies in mouse^6,13,24–26^ and human embryos^6^, have found differences in mitochondrial physiology and activity between cells in the TE and ICM.

During an in vitro fertilization (IVF) cycle, exogenous hormones are administered to a patient in order to stimulate the ovaries to produce multiple oocytes and subsequent embryos. Developing tools to select the best embryo to transfer has been a long term focus in IVF^27–29^. Improved embryo selection would decrease the number of transfers a patient must undergo, which would in turn, reduce its economical and emotional cost^21,30^. Current selection methods primarily rely on embryo morphology assessments^31^. Blastocyst morphology grading is typically based on the expansion stage, the consistency of both the ICM and TE, and the time to blastocyst formation since fertilization^31^. Despite its widespread use, embryo morphological assessments have limited predictive power, and the further disadvantage that they are unable to provide a direct measure of the physiology of the embryo^32^. Alternative embryo selection approaches include: invasive preimplantation genetic testing for aneuploidy (PGT-A), which is increasingly being used^33–35^; time-lapse imaging^36^, which provides detailed information on developmental dynamics; and, more recently, artificial intelligence^37^. Additionally, several studies in mouse and human embryos showed a clear association between metabolic function and implantation potential^5,13,21,38^. Embryo morphology^39^ and time to blastocyst formation^40^ have also been found to be linked with embryo metabolism. This suggests that measures of metabolism might provide a means to select high quality embryos for transfer, but approaches based on this premise have so far not been successful^40,41^.

Many approaches to study embryo metabolism hinge on either intracellular measurements^42,43^ or quantification of metabolites in the spent media to detect the metabolic activity of the whole embryo^5,21,28,44,45^. These methods are often either invasive or require highly specialized skills to perform. Recently, non-invasive methods to measure intracellular metabolic activity of oocytes and embryos have also been pursued. Several endogenous molecules like nicotinamide adenine phosphate dinucleotide (NADPH), nicotinamide adenine dinucleotide (NADH) and flavine adenine dinucleotide (FAD+) are autofluorescent^46^. These molecules are electron carriers that play key roles in metabolic pathways, which makes them ideal candidates for characterizing cellular metabolism^47,48^. The fluorescence spectra of NADH and NADPH are almost indistinguishable^49^, therefore the combined fluorescence of NADH and NADPH is often referred to as the NAD(P)H signal. NAD(P)H^50,51^ and FAD+^22,52^ autofluorescent intensity can be used to measure switches in the metabolic state of cells throughout development.

Additional information on NAD(P)H and FAD+ can be obtained using fluorescence lifetime imaging microscopy (FLIM)^53^. FLIM not only provides information on fluorescence intensity, but also gives information on the fluorescence lifetime, i.e. the time the fluorophores remain in their excited state. The fluorescence lifetimes of NAD(P)H and FAD+ depend on their microenvironment, including their engagement with enzymes, and thus provides a sensitive means to characterize variations in metabolic state^49^. FLIM enables non-invasive, quantitative measurements of the metabolic state of mouse embryos^54–56^, but its application to human preimplantation embryos has yet to be established^57^. FLIM can be performed with two-photon fluorescence microscopy^53,58^, which allows intrinsic sectioning and deep imaging^59^. Another advantage of performing FLIM with two-photon microscopy, is that it enables simultaneous non-invasive imaging of spindle morphology via second harmonic generation (SHG). SHG is a non-linear phenomenon that occurs when light scatters from highly ordered structures that lack inversion symmetry, like the spindle^54,59–61^. Non-invasive techniques to simultaneously measure the cellular metabolic state and spindle dynamics of human embryos will offer a better understanding of basic embryo biology and may assist in improving embryo screening in a clinical setting.

In the present study, we used non-invasive FLIM to measure the metabolic state of human blastocysts. We explored the extent to which FLIM can quantitatively measure metabolic changes through human blastocyst expansion and hatching. We studied spatial patterns in the metabolic state within human blastocysts and their association with stage of expansion, day of development since fertilization and morphology. Finally, we explored the sensitivity of this technique in detecting metabolic variations between blastocysts from the same patient and between patients.

## Results

### Two-photon microscopy of endogenous autofluorescence and second harmonic generation imaging enable visualization of subcellular structures in preimplantation human embryos

Preimplantation human embryos are often visualized using bright-field microscopy in IVF clinics, which is sufficient for blastocyst morphological grading but provides limited cellular and subcellular information (Figure 1A). We first investigated what additional morphological information can be provided by two-photon microscopy of endogenous NAD(P)H and FAD+. Two-photon microscopy enables deep tissue imaging with optical sectioning^59^, allowing us to perform three dimensional (3D) reconstruction of blastocysts by combing multiple Z plane images of NAD(P)H autofluorescence (Figure 1B). Cell nuclei appear as dark ovals in these 3D reconstructions (Figure 1B, arrow), as confirmed by comparison to 3D reconstruction of human embryos stained for DNA with Hoechst (Figure 1C). As FAD+ is significantly more enriched in mitochondria than NAD(P)H in many systems^47,48^, we sought to determine the extent to which two-photon microscopy of FAD+ can provide information on mitochondrial localization in human preimplantation embryos. We stained five human blastocysts with MitoTraker Red CMXRos, a dye that specifically labels mitochondria, and simultaneously imaged MitoTraker and FAD+ autofluorescence (Figure 1D). We used machine-learning based software (Illastik, version 1.0^62^) to segment bright intracellular regions in both the MitoTraker and FAD+ images and found an overlap of photons between regions of 89±8%. Hence, the overwhelming majority of the FAD+ signal in human blastocyst is associated with mitochondria, and thus FAD+ imaging provides information on mitochondrial localization. The same laser illumination used for two-photon microscopy can be simultaneously employed for SHG, which we combined with FAD+ imaging. Using both autofluorescence and SHG imaging, it was possible to observe spindles in mitotic cells in human blastocysts, along with the subsequent cell division (Figure 1E). Thus, two-photon microscopy of NAD(P)H and FAD+, combined with SHG imaging, provides a non-invasive means to image cellular and subcellular structures in human blastocysts, including mitochondria, nuclei and spindles.

**Figure 1.**
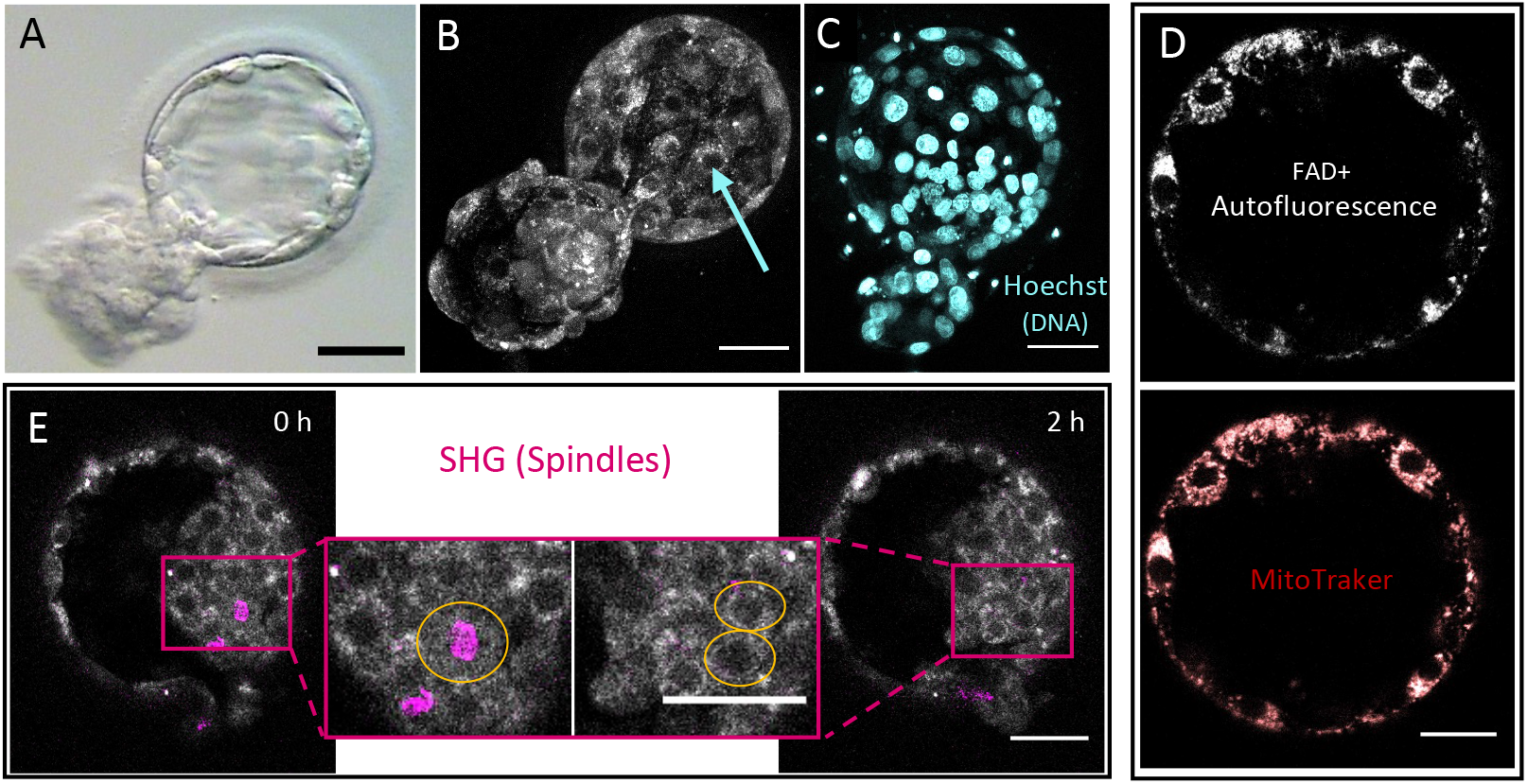
Two-photon fluorescence lifetime imaging microscopy (FLIM) and Second Harmonic Generation (SHG) enable visualization of cellular and subcellular structures. (A) Standard bright-field image of a human discarded blastocyst. (B) 3D reconstruction of two-photon NAD(P)H intensity images from multiple focal planes of the same blastocyst. Deep tissue imaging using two-photon microscopy enables the detection of subcellular structures such as the nucleus (blue arrow). (C) 3-D reconstruction of DNA staining Hoechst showing cell nuclei. (D) Simultaneous FLIM FAD+ autofluorescence of a human blastocyst and mitochondria dye MitoTraker Red CMXRos demonstrate a high colocalization. (E) SHG imaging, in combination with FLIM NAD(P)H autofluorescence imaging enables detection of spindle formation capturing cellular mitotic divisions (E). Scale bars, 40 μm.

### Metabolic variations during human blastocyst expansion and hatching

To study the changes in metabolism during the expansion and hatching of preimplantation embryos, we performed time-lapse imaging of NAD(P)H (Figure 2A), FAD+ (Figure 2B, grey) and SHG (Figure 1B, magenta) (Movie S1) of 10 morphologically normal human blastocysts. We acquired three Z planes for each channel every two hours, over a 36-hour period. All blastocysts progressed through development, as assessed by a senior embryologist. Hence, this imaging did not appear to disrupt human embryonic preimplantation development.

**Figure 2.**
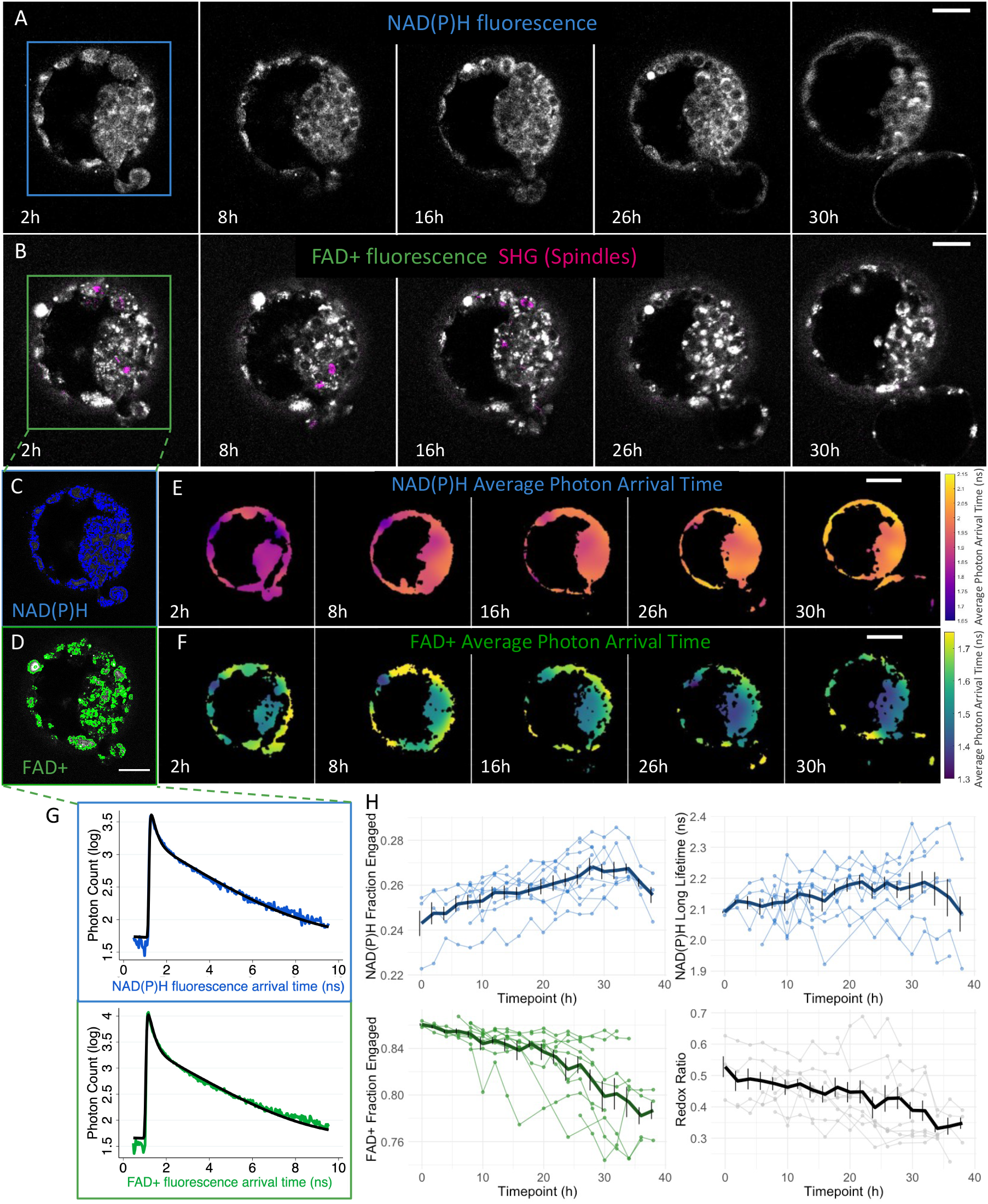
Non-invasive FLIM and SHG imaging detect metabolic variations during human blastocyst development. FLIM time-lapse imaging of the autofluorescence of NAD(P)H (A), and (B) of FAD+ (in grey) and SHG spindle imaging (in magenta) of a human blastocyst throughout 36h of incubation. Supervised machine learning was applied to create masks to segment the fluorescence signal of NAD(P)H (C) or FAD+ (D). (E) FLIM time-lapse imaging of the photon arrival times of NAD(P)H and (F) of FAD+ enable visualization of variations of metabolic state of human blastocysts throughout expansion and hatching. Color bars show the average photon arrival times in nanoseconds (Movie S1). (G) All photon arrival times from a single embryo mask were combined to create a fluorescence decay for each fluorophore (NAD(P)H in blue and FAD+ in green). These curves were fitted to a two-exponential model (black curves) to the quantitative parameters for characterizing the metabolic state of an embryo. (H) Time trajectory plots of individual embryos (n=10) of four of the metabolic parameters produced: NAD(P)H fraction engaged, NAD(P)H long lifetime, FAD+ fraction engaged and redox ratio (SI1). The average time curves from all the embryos are shown in thicker lines with SE bars, showing that these trends were reproducible among blastocysts. Scale bars, 40 μm.

We used an intensity-based machine learning algorithm (see methods) to segment both the NAD(P)H (Figure 2C) and the FAD+ signal (Figure 2D). As an initial means to investigate the metabolism of human blastocysts, we calculated the average photon arrival time (methods) throughout the segmented regions for both NAD(P)H (Figure 2E) and FAD+ (Figure 2F). Both signals showed complex spatial patterns which evolved over time as the embryos progressed through development. To better quantify these temporal changes, we grouped all photons from the segmented regions to obtain histograms of NAD(P)H (Figure 2G, upper) and FAD+ (Figure 2G, lower) photon arrival times, which were fit to two-exponential decay models (Figure 2G, black lines). This provided nine metabolic FLIM related parameters at each point in time: fluorescence intensity, short and long lifetimes, and the fraction of molecules engaged with enzymes for both NAD(P)H and FAD+, as well as the redox ratio (NAD(P)H intensity/ FAD+ intensity). These FLIM parameters showed stereotypical changes over time during blastocyst development, with some parameters increasing and some decreasing as the embryos progressed past the early blastocyst stage (Figure 2H and SI1).

### Spatial variation in human blastocysts metabolic parameters

Continual time-lapse microscopy, as described above, provides detailed information on the evolution of individual human blastocysts, but presents practical limits on the number of embryos that can be imaged. We thus obtained additional FLIM data on human blastocysts at single time points, acquired two hours after thawing. We imaged a total of 215 human blastocysts from 137 patients that were discarded and donated for research.

We first used this data to further investigate the spatial variation in metabolic parameters in human blastocysts. We segmented the images and visualized the spatial distribution of average photon arrival times for NAD(P)H (Figure 3A) and FAD+ (Figure 3B), which appeared to indicate different metabolic signatures in the ICM and TE. These spatial patterns disappear after randomizing the photon arrival times for each pixel in the segmented region (Figure SI 2), indicating that the observed spatial patterns are real and not an artifact of the averaging procedure or the geometry of the blastocyst (Methods). To further investigate the difference in metabolism between the ICM and TE, we manually segmented them and grouped photons from each region to determine their FLIM parameters (n=187). All metabolic parameters displayed significant differences between ICM and TE as measured by a paired t-test with post-hoc correction using Benjamini - Hochberg’s false discovery rate (FDR), (FDR *p*<0.0001). We were concerned that the different number of photons acquired from the two regions might artificially contribute to the measured difference in FLIM parameters. We therefore repeated the analysis, but by randomly selecting the same number of photons from the ICM and the TE. All the significant differences were upheld (FDR *p*<0.0005), indicating that the measured difference in FLIM parameters is robust to the number of photons acquired. The observed differences in FLIM parameters between ICM and TE argues that cells in these regions are in different metabolic states (Figure 3C). To further investigate this on the single embryo level, we calculated the ratio of each FLIM parameter between the ICM and TE of every embryo. While the average of each ratio across embryos was significantly different from 1 (FDR *p*<0.0001), there was substantial embryo-to-embryo variability (Figure 3D).

**Figure 3.**
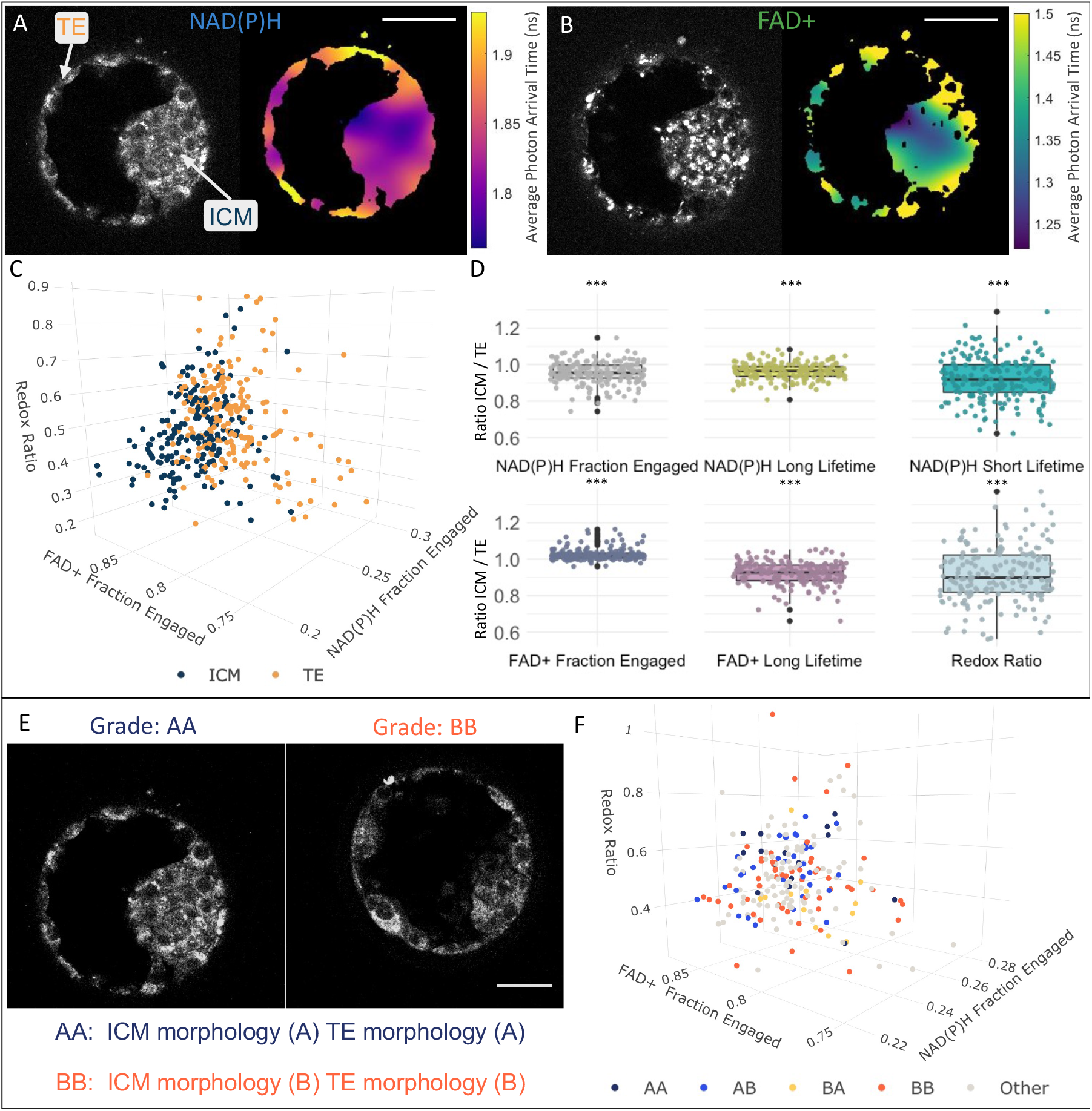
Variations in metabolic signatures between inner cell mass (ICM) and trophectoderm (TE) of human blastocysts. FLIM intensity image (left) and image of the average photon arrival time (right) of NAD(P)H (A) and of FAD+ (B) reveal spatial pattern of average photon arrival times, suggesting different metabolic state between the ICM and TE. Color bars show the average photon arrival times in nanoseconds. (C) FLIM parameters with the largest separation between ICM (blue) and TE (orange) were plotted in this 3D plot (*FDR p* <0.001). Each dot corresponds to the mean of 3Z images of the ICM or TE of single embryo. (D) Ratios of the FLIM parameters between the ICM and the TE. The box plots depict the interquartile range of the ratio, the center of the box represents the mean, the horizontal line in the box represents the median, and the vertical lines represent the 5 and 95% quartiles. Black dots represent points outside of the interquartile range. Color dots represent the average ratio per each individual embryo. ICM and TE morphology ^31^ of each embryo at the time of imaging. (E) Example of NAD(P)H intensity images of two human discarded blastocysts with varying morphological grading (AA and BB). (F) FLIM parameters were compared between blastocysts of different morphological categories (AA, AB, BA, BB, and other), this 3D plot shows no significant differences of metabolic signatures between embryo morphology grades (*FDR p* >0.05). Each dot corresponds to the mean of an embryo. Scale bars, 40 μm. *** signifies *FDR p*< *0.001*.

We next explored if the embryo-to-embryo differences in FLIM of TE and ICM was related to the known large variations in TE and ICM morphology in human blastocysts. A senior embryologist graded the ICM and TE morphology of each embryo at the time of imaging using the standard grading system described by Schoolcraft et al. ^31^. Briefly, an ICM with many, compacted cells are considered to be grade A, if it has several cells and they are loosely grouped, it is grade B, whereas if it contains very few cells, it is considered grade C. For the TE, a score A is assigned if the blastocyst has many cells that form a cohesive epithelium, if it has fewer cells, then it is graded as B, and if the TE has very few, large cells it is given a grade of C. These embryos then were divided into five categories: 16 embryos with grade AA, 33 with grade AB, 35 with BA, 12 with BB and 99 were categorized as other (any with C). The first and second letter corresponds to the ICM and TE grade, respectively. None of the FLIM parameters showed significant differences between morphological groups (FDR *p*>0.05) (Figure 3F). Thus, the embryo-to-embryo differences in FLIM of TE and ICM is not associated with variations in TE and ICM morphology.

### Temporal variations of human blastocysts metabolic state

#### Blastocyst stage

To further study the metabolic shifts observed during blastocyst development, we explored the relationship between FLIM parameters, averaged over the entire embryo, and embryo stage. A senior embryologist assigned each human discarded blastocyst to one of four stages (Figure 4A): early blastocysts (N = 25 embryos), expanded blastocyst (N = 58), hatching blastocyst (N = 101), and fully hatched blastocyst (N = 31). We computed the percentage relative to early blastocyst for each FLIM parameter and stage. FAD+ fraction engaged, FAD+ long lifetime, FAD+ short lifetime, and redox ratio all exhibited significant variations (FDR *p*<0.001), with the largest occurring for hatching and fully hatched blastocysts (Figure 4B). We next investigated the potential existence of subtle changes in FLIM parameters during blastocysts expansion by testing for correlations between the diameter of early and expanding blastocysts with FLIM parameters. There were no significant correlations between these metabolic profiles and the diameter of the blastocysts (FDR *p*>0.05). Thus, the largest changes in FLIM parameters are associated with the process of blastocyst hatching.

**Figure 4.**
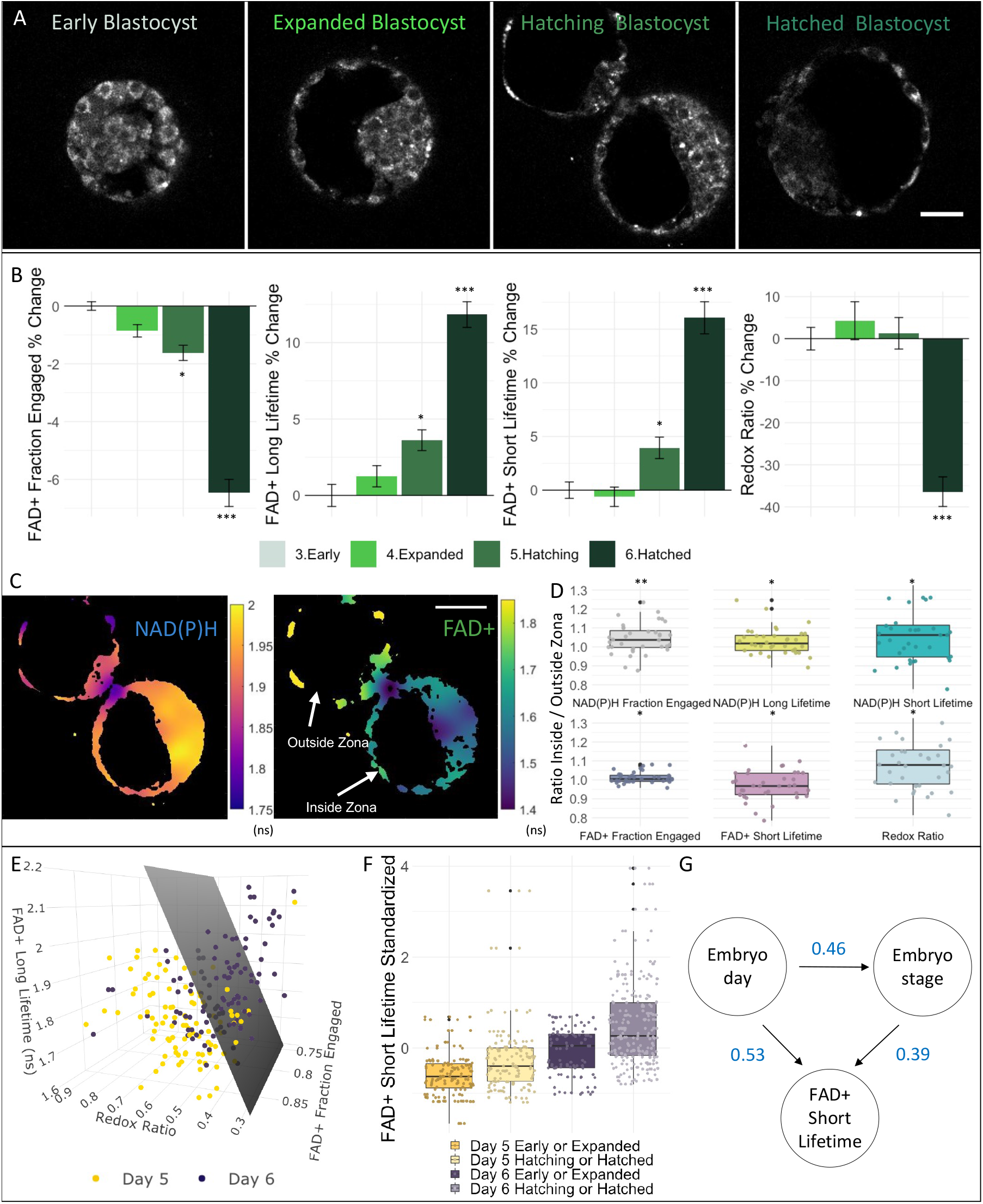
Human blastocyst metabolic state is associated with embryo day and embryo expansion stage. (A) NAD(P)H intensity images of four human blastocysts with varying expansion grade: early, expanded, hatching and hatched blastocysts. (B) Bar plots of the percentage change from early blastocysts of FAD+ fraction engaged, FAD+ long lifetime, FAD+ short lifetime and redox ratio with standard error bars displayed, showing a distinctive variation in these FLIM parameters between early and hatched blastocysts. (C) FLIM image of the photon arrival times (right) of NAD(P)H and of FAD+ (left) showing variations in the metabolic state between the portion of the TE inside and outside the zona pellucida. Color bars show the average photon arrival times in nanoseconds. (D) Of those embryos that were hatching (n=36), we computed the ratio of the FLIM parameters of the TE between the inside and outside of the zona pellucida. The box plots show the interquartile range of the ratio, the horizontal line in the box represents the median, and the vertical lines represent the 5 and 95% quartiles. Black dots represent data points outside of the interquartile range. Color dots represent the average ratio of each blastocyst. (E) FLIM parameters with largest separation between embryos of either day 5 or 6 after fertilization were plotted in a 3D plot. Each dot corresponds to the mean of a single blastocyst. Support vector machine was used to create a hyperplane that best separates day 5 and day 6 blastocysts (grey plane). (F) We subdivided the data in four categories: day 5 early/expanded, day 5 hatching/hatched, day 6 early/expanded or day 6 hatching/hatched embryos. This box plot displays the standardized FLIM FAD+ short lifetime of these four categories, showing that both day and stage impact FAD+ short lifetime. (G) DAG of FAD+ short lifetime showing that embryo day and stage are correlated and that embryo FAD+ short lifetime is dependent both on embryo day and stage. Numbers in blue represent the coefficient of the multilevel model. Scale bars, 40 μm. *** signifies *FDR p* <*0.001*, \*\**FDR p* <*0.01* * signifies *FDR p* <*0.05*.

To better understand the metabolic variations associated with hatching, we examined the differences in FLIM parameters between the portions of TE inside the zona pellucida compared to those outside. We analysed 36 hatching blastocysts from our cohort that had a substantial area both inside and outside the zona pellucida (Figure 4C). We computed the ratio of FLIM parameters between the TE portion of the blastocyst inside and outside the zona pellucida. NAD(P)H intensity (FDR *p*<0.02), fraction engaged (FDR *p*<0.02), NAD(P)H long (FDR *p*<0.03) and short lifetime (FDR *p*<0.03), FAD+ fraction engaged (FDR *p*=0.03), FAD+ short lifetime (FDR *p*=0.04), and the redox ratio (FDR *p*=0.02), all exhibit significant differences between the TE portion of the blastocyst inside and outside the zona pellucida (Figure 4D).

#### Day of development since fertilization

The discarded human blastocysts we studied were vitrified at different periods of time after fertilization, the majority either on day 5 (N = 98 embryos) or on day 6 (N = 111). We used multilevel models to compare FLIM parameters of day 5 and day 6 blastocysts. We found significant variations in NAD(P)H intensity (FDR *p*=0.002), FAD+ intensity (FDR *p*=0.01), FAD+ fraction engaged, FAD+ long and short lifetime, and redox ratio (FDR *p*<0.0002). The separation was so strong that a support vector machine generated hyperplane based on just three FLIM parameters gave predictions of whether an embryo was day 5 or day 6 with an accuracy of 77% (Figure 4E).

Embryos develop to later stages over time, so, it would be possible that differences in FLIM parameters between day 5 and day 6 embryos might be due to embryos being at different developmental stages on these days. To investigate this, we explored the conditional dependencies between the day of embryo development since fertilization, stage of expansion and the FLIM parameters that were significantly associated with both of those (FAD+ fraction engaged, FAD+ long lifetime, FAD+ short lifetime and redox ratio). We began by investigating FAD+ short lifetime. Subdividing the embryos into two different stage categories (early/expanded or hatching/hatched) on both day 5 and day 6 demonstrates that day and stage both impact FAD+ short lifetime (Figure 4F). Upon conditioning on stage of expansion, the partial correlation between day of embryo development and FAD+ short lifetime, (day, FAD+ short lifetime | stage), is 0.53 (p<0.0001). Upon condition on day of embryo development, the partial correlation between embryo stage of expansion and FAD+ short lifetime, (stage, FAD+ short lifetime| day), is 0.39 (p<0.0001). Thus, expansion stage and day of embryo development are both associated with FAD+ short lifetime. Furthermore, day of embryo development and expansion stage are correlated with a coefficient of 0.46 (*p*<0.00001), after conditioning on FLIM parameters. These conditional dependencies can be represented by a directed acyclic graph (DAG), which shows that day of embryo development, expansion stage and FAD+ short lifetime all depend on each other (Figure 4G). Similar results held for the other FLIM parameters (Figure SI3).

### Metabolic variance between human blastocysts, between patients and between images

Embryo day of development and expansion stage are two factors which lead to differences in metabolism between different human preimplantation embryos. There are presumably other factors that also contributed to these differences. Some of these factors may cause differences in metabolism between embryos from different patients, whereas other factors may cause differences in metabolism between different embryos from the same patient. Indeed, plotting FLIM measurements from different embryos from the same patient often leads to a clear separation between embryos (Figure 5A). In contrast, plotting FLIM parameters from embryos from different patients often produces substantial overlap (Figure 5B). Thus, factors that produce differences in metabolism between embryos from different patients appear to be more subtle than the factors that produce variations between embryos from one patient. To more systematically quantify these effects, we used multilevel models ^63,64^ to determine the residual variance in FLIM parameters explained by three levels: the differences between patients, between embryos from the same patient, and between images taken at different Z positions of the same embryo (Figure 5C). The residual variances of FLIM parameters for each level were significantly different than 0 (FDR p>0.05), except the variance of NADH Long Lifetime and FAD+ fraction engaged between patients. For the fluorescence lifetimes, the fractions engaged, and the redox ratio, the variance associated with differences between patients and between different Z positions were smaller than the variances associated with differences between embryos from the same patient.

**Figure 5.**
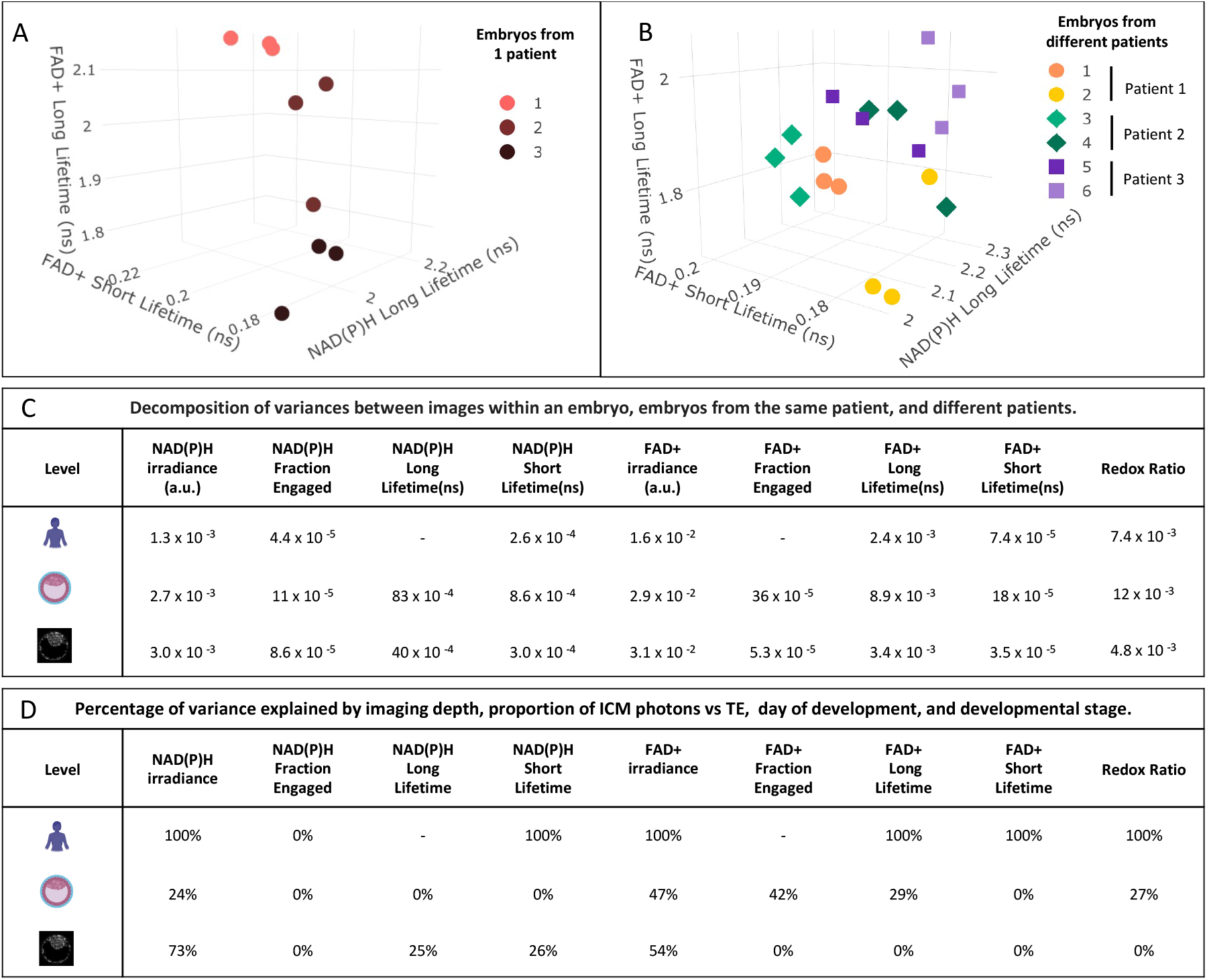
Variance of the metabolic parameters between patients, embryos, and images. Multilevel models encode information on the variance of each FLIM parameter between different patients, between different embryos from the same patient, and between different images from the same embryo (taken at different Z positions). (A) Example of a patient with three different blastocysts. Colors correspond to a different embryo and each dot corresponds to a different image, from a different Z position, within the embryo. (B) Example of three patients with two embryos each imaged. Symbols correspond to different patients; each color corresponds to a different embryo and each dot corresponds to a different Z position within each embryo. (C) The variance of each FLIM parameter associated with each level: patient (top), embryo (middle) and images (bottom). FLIM parameter variances between patients were significantly different from 0 (FDR p values ranging between 0.001 and 0.04), except NADH long lifetime and FAD+ fraction engaged. All variances associated with embryo level (FDR p values ranging between 1.1 x 10^−6^ and 9 x 10^−9^) and all variances between images within embryos were significantly different from 0 (p values < 1 x 10^−10^). (D) The percentage of variance of each FLIM parameter associated with each level explained after controlling for embryo day, expansion stage, imaging depth and the proportion of photons coming from the ICM vs TE. 100% corresponds to no statistically significant variance remaining after controlling for the listed factors. 0% corresponds to the variances not being altered after controlling for the listed factors.

Finally, we investigated factors that might lead to the differences in FLIM parameters associated with these different levels (Figure 5D). Over 70% of the variance in NAD(P)H intensity and over 50% of the variance in FAD+ intensity between different Z positions of the same embryo can be explained by a combination of imaging depth and the proportion of photons coming from the ICM vs TE. Surprisingly, we found no association between FLIM parameters of blastocysts and maternal age at oocyte retrieval, patient body mass index (BMI), or levels of anti-Mullerian hormone (FDR p>0.05), all of which are known to be associated with successful IVF outcome^31,65,66^. Differences in FLIM parameters between patients were largely explained by differences in day of embryo development and expansion stage: after controlling for these two factors, NAD(P)H fraction engaged was the only FLIM parameter with statistically significant variance at the patient level. In contrast, there was still substantial variance in FLIM parameters between embryos from the same patient, even after controlling for day of embryo development, expansion stage, imaging depth, and the proportion of photons coming from the ICM vs TE. Thus, there are metabolic difference between embryos from the same patient which cannot be accounted for by these factors.

## Discussion

In this study, we explored the use of FLIM of NAD(P)H and FAD+ for measuring the metabolic state of human blastocysts. We found that the metabolic state of blastocysts changes continuously over time and varies based on both time since fertilization and developmental stage. We also observed distinct metabolic states between cells in the ICM and the TE, and between cells inside and outside the zona pellucida. Finally, we demonstrated that FLIM is sensitive enough to detect metabolic variations in individual blastocysts from the same patient and variations between patients.

Impaired metabolic activity results in decreased developmental capacity in oocytes and embryos, both in mouse^4,14^ and human^4,15^. To this end, several reports have focused on detecting metabolic biomarkers of embryo quality^4–6,16,21,45,46,50^. In this study, we have demonstrated the use of FLIM imaging to detect the metabolic state of human discarded blastocysts. We observed that combining FLIM with SHG imaging^60,61,68^ provides information on a range of cellular and subcellular structures in human blastocysts, including mitochondrial localization and physiology, spindle dynamics and mitotic divisions. Taken together, these results demonstrate that FLIM and SHG imaging can be used to obtain quantitative, non-invasive and biologically relevant information on live human blastocysts.

Metabolic shifts from the morula to the blastocyst stage have been described previously^1–3,41^, including changes in redox state, and increasing levels of glucose uptake, oxygen consumption and lactate production^10–12,69^. Our results extend these prior reports by showing continuous variations in metabolic state during human blastocyst development from the early blastocyst to hatching stages. Thus, in contrast with previous reports^70^, we observed highly significantly different metabolic states in human blastocysts at day 5 vs day 6 post fertilization. Human embryos develop at different rates, and our results show that only a fraction of the metabolic differences between blastocysts at different days post fertilization can be explained by embryos tending to be at later developmental stages at later times. This implies an association between embryo developmental rate and metabolism, as has also been seen by other approaches^5,18,52^ Taken together, these observations argue that the metabolism of human blastocysts are highly dynamic and flexible. It is an exciting future challenge to unravel the mechanistic basis of these metabolic variations in human embryos.

Our measurements showing different metabolic state of cells in the ICM and TE are consistent with prior work indicating that they contain distinct mitochondrial physiology and function^6,24–26^. Thus, FLIM is sensitive enough to detect physiologically relevant changes in metabolism in human blastocysts, as has been previously seen for differentiation in *Caenorhabditis elegans*^71^ and mouse neuronal development^72^. We also observed differences in the metabolic state of trophectoderm cells inside and outside of the zona pellucida during hatching. To our knowledge, this has not been reported previously. There seem to be two possibilities: 1) these metabolic differences may reflect differing microenvironments. Perhaps the increase in lactate production, and associated pH change, during hatching^10^ reflects the different energy expenditure needed by TE cells inside and outside the zona pellucida, 2) the hatching process itself may also lead to metabolic changes, either due to specific signaling pathways or a general response to associated stresses^16^.

In addition, by analyzing our data using multilevel modeling^63,64^, we found that blastocysts from the same patient have significantly different metabolic states. While some of this variation can be accounted for by time post fertilization and stage of development, a substantial variance between embryos remains even after accounting for these factors. It is unclear what causes these additional variations in metabolism between embryos. One possibility is that metabolism adjusts to respond to cellular or DNA damage, which are prevalent in preimplantation human embryos^18,19,23^. This provides an attractive explanation for the previously observed associations between blastocyst metabolism and viability^5,13,21^. However, while embryo morphological scores are also associated with viability^31,44,73,74^, we found no association between embryo metabolic state and morphological scores. This implies that multiple distinct processes impact embryo viability, some of which are associated with metabolism and others of which are associated with morphology. It will be of both fundamental interest and potential clinical import for future work to disentangle the contribution of specific pathways to embryo viability. Furthermore, the lack of association between metabolic state and morphology scores suggests that FLIM might provide synergistic information with conventional microscopy to aid in selecting the highest quality embryo for transfer in Assisted Reproductive Technologies (ART).

There are also limitations to this study. First, the human blastocyst analysed were vitrified, thawed, and warmed for two hours prior to FLIM imaging. Whether vitrification causes metabolic distress in human blastocysts remains unknown, even though its clinical success has been well documented^75^. In addition, it does not seem to alter mitochondrial potential or intracellular reactive oxygen species^76^. Additionally, FLIM imaging did not disrupt normal blastocyst development nor has it impacted mouse embryo viability^56^. These observations further support the potential use and the safety of FLIM as a non-invasive technique. Second, the blastocysts used in this study were discarded material, and include both euploid and aneuploid embryos. Our preliminary analysis indicates some associations between embryo metabolic state and ploidy^77^, which will be reported on in a subsequent manuscript. Future studies should focus on addressing this important subject in more detail. Surprisingly, we did not observe a correlation between blastocyst metabolic state and maternal age or BMI, each of which have long been linked with embryo quality^31,65^ and mitochondrial function^78,79^. This result must be interpreted with caution as it may reflect a selection bias in the material used in this study: we only imaged embryos that were discarded, yet also reached the blastocyst stage. A clinical study using non-discarded embryos of all preimplantation developmental stages will provide a more complete view the effect of maternal characteristics on embryo metabolic state. Finally, recent work has established a framework to relate FLIM parameters in oocytes to mitochondrial metabolic fluxes^80^, but additional work is required to determine if this is applicable to the later stage embryos studied here.

In summary, non-invasive metabolic imaging via FLIM is a promising tool for quantitatively characterizing the metabolic state of human blastocysts. FLIM can sensitively detected metabolic variations associated with blastocyst development over 36h, the time post fertilization and blastocyst developmental stage. FLIM revealed differences in metabolic state between cells in the ICM and the TE, and between TE cells inside and outside the zona pellucida. The lack of an association between the metabolic state of human blastocyst and their morphological grading indicates that these two assessments may provide synergistic information to improve blastocyst selection. However, future work is required to determine the extent to which FLIM measurements are associated with the probability that a transferred human blastocyst will lead to a live birth.

## Materials and Methods

### Sample preparation

Human blastocysts were discarded and donated for research under determinations by the Beth Israel Deaconess Medical Center and New England institutional review boards (New England IRB WO 1-6450-1). Two hundred and fifteen vitrified human blastocysts from 137 patients, with a mean ± standard deviation age of 35.4 ± 4.7 years, BMI of 25.9 ±5.2 kg/m^2^, and mean AMH of 3.55 ± 2.9 ng/mL were analyzed in this experiment. These embryos were thawed according to the manufacturer’s recommendations (90137-SO - Vit Kit-Thaw, FUJIFILM Irvine Scientific, USA) and cultured for two hours in individual drops of 50μl of Continuous Single Culture Complete (CSC) media with human serum albumin media (HSA) (FUJIFILM Irvine Scientific, USA) overlain with mineral oil in an incubator at 37C, 7% CO_2_ and 6% O_2_. Patient clinical characteristics such as age, BMI and AMH hormone levels were provided in a de-identified database. Blastocyst’s morphological grade^31^ were evaluated at the time of imaging by a senior embryologist. An ICM and TE grade A, B, or C was assigned for each blastocyst. Stage of development (early, expanded, hatching or hatched blastocyst) and day since fertilization (day 5 or 6) was also monitored for each embryo.

### Imaging Protocols

Discarded blastocysts were transferred and imaged in a custom glass-bottomed microwell dish with 80uL of media overlain with mineral oil, in an on-stage incubation system (Ibidi GmbH, Martinsried, Germany) to maintain environmental culture conditions. For the blastocyst development time-lapse experiments, metabolic images were taken at three Z-planes 7 um apart every two hours over the course of 36 h, using 12mW for NAD(P)H and 20mW for FAD+ and 60 seconds of integration time for each plane. For the blastocyst experiments, metabolic images were taken once at three Z-planes 7 um apart, using 12mW for NAD(P)H and 20mW for FAD+ and 60 seconds of integration time for each plane.

### Staining protocols

For MitoTraker experiments, five blastocysts were incubated with 5nM MitoTraker Red CMXRos (M7512, Thermo Fisher, USA) for 20 minutes. For DNA staining experiments, embryos were stained with 1ug/mL of Hoechst (Thermo Fisher, USA) for 20 minutes. The samples were then washed three times in CSC + HSA and transferred to a glass bottom dish for imaging.

### Fluorescence Lifetime Imaging Microscopy (FLIM)

FLIM measurements were performed on a Nikon TE300 (Nikon, Japan) microscope using two-photon excitation from a Ti:Sapphire pulsed laser (M-squared Lasers, UK) with a 80MHz repetition rate and 150 fs pulse width, a Galvano scanner, TCSPC module (SPC-150, Becker and Hickl, Germany) and a hybrid single photon counting detector (HPM-100-40, Becker and Hickl, Germany). Imaging was performed with a 20X Nikon objective with 0.75 numerical aperture (CFI Apo 20X, NA 0.75, Nikon). The wavelengths of NAD(P)H and FAD+ excitation were set to 750nm and 890nm, respectively. The powers measured at the objective used were 12mW for NAD(P)H and 20mW for FAD+. Optical bandpass filters were positioned in a filter wheel in front of the detector – 447/60 nm for NAD(P)H (BrightLine, Semrock, USA) and 550/88nm for FAD+ (Chroma technologies) with an additional 650nm short pass filter mounted on the detector (Chroma technologies). SHG was detected simultaneous with FAD imaging by a single-photon counting detector (PMC-150, Becker-Hickl GmbH, Germany) in the forward direction, with 650 short-pass and 440/20-nm bandpass filters (BrightLine, Semrock, USA). Each NAD(P)H and FAD+ image was acquired with 60 seconds of integration time. A customized motorized stage (using CONEX TRA12CC actuators, Newport, USA) was used to perform multi-dimensional acquisitions. We acquired three FLIM images varying Z axis per each human blastocyst. All the electronics were controlled by SPCM software (Becker and Hickl, Germany) and custom LabVIEW software.

### Data Analysis

Data was analyzed using a customized MATLAB version R2019b (MathWorks, USA) code. Samples were incorporated to the analysis according to the information available. NAD(P)H and FAD+ images were trained using a supervised machine learning segmentation software (Illastik, version 1.0^62^) to classify pixels in intensity images into either NAD(P)H or FAD+ signal from the cells or the background. The algorithm was trained using 40 random NAD(P)H and FAD+ intensity images. For each cell segment, the photon arrival time histogram was modeled as a bi-exponential decay:

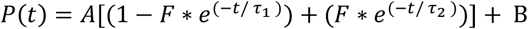

Where, A is a normalization factor, B is the background, τ_1_ is the short lifetime, τ_2_ is the long lifetime and *F* is the fraction of molecule with long lifetime (fraction engaged with enzymes for NAD(P)H and unengaged for FAD+). This function was convolved with a measured instrument response function to model the experimental data, and the least square fitting yielded quantitative values for these fit parameters. The fluorescence intensity was calculated for each embryo dividing the number of photons by the area of the embryo. An additional parameter, the redox ratio (NAD(P)H fluorescence intensity/FAD+ fluorescence intensity) was also calculated per image. Together, a single FLIM measurement produced 9 parameters, 4 for NAD(P)H and 4 for FAD+ and redox ratio, to quantitatively characterize the metabolic state of human blastocysts.

The average photon arrival time of NAD(P)H and FAD+ is computed for each pixel within the embryo by averaging over the directly measured photon arrival time of all photons coming from NAD(P)H and FAD+ within a single pixel. Since the single pixel average is noisy due to limited photon counts per pixel, we smoothened the data by averaging each pixel over its neighboring pixels weighted by a gaussian kernel depending on the distance between the pixels. The standard deviation of the gaussian kernel is 20 pixels. We only averaged over pixels within the embryo and exclude pixels of the background and the cavity of the embryo. As a statistical control, we randomized the photon arrival times in each pixel by drawing a random photon arrival time from a gaussian distribution with a mean of the average photon arrive time and a variance equal to that of the photon arrival time distribution of the blastocyst. This procedure produces a randomized photon arrival time image of the blastocyst with the exact same geometry. We repeated the average photon arrival time calculation described above for this “control image” to test if there is any artifact associated with the averaging procedure. The control image consistently produced a homogeneous average photon arrival time, suggesting the heterogeneity observed in the blastocyst is of a biological origin rather than statistical artifact.

### Statistical Analysis

All statistical tests were performed using Stata Statistical Software version 16.0 (LLC Stata Corp, Texas, USA) and R Studio Version 1.3.959 (R Foundation for Statistical Computing, Vienna, Austria). Our data was structured hierarchically, three images per embryo (i) and 1 – 7 embryos (j) per patient (k). Therefore, we used multilevel models with restricted maximum likelihood estimates^63^ to analyze this structured data. We incorporated the corresponding predictors (embryo morphology, day, expansion stage or patient age, BMI and AMH) for each analysis, using the multilevel model:

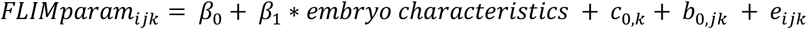

Where β_1_ corresponds to the intercept; α_1_ to the slope; *embryo characteristics* to the day, stage or morphological grading of the embryos; *c*_0,*k*_ is the patient level random error; *b*_0,*jk*_ is the embryo level random error; *e*_*ijk*_ is the image random error^63^. This modelling encodes information on the variance associated with each level: patient, embryos within patients and images within an embryo. One tailed Z-test was performed to determine whether these variances were significantly different than zero.

Additionally, paired t-test were performed to detect metabolic variations between the ICM and the TE of each embryo. For all comparisons of ratios, one sample t-test was performed to determine whether these ratios were significantly different than one. Furthermore, we calculated the percentage change of all metabolic parameters between each expansion stage compared to the early blastocyst stage and pairwise comparisons were performed to analyze their significance. All p-values were corrected for multiple comparisons using Benjamini - Hochberg’s false discovery rate (FDR), at a *q* value of 0.05. FDR *p*-values of <0.05 were considered statistically significant.

We used a support vector machine algorithm (SVM)^81^ to fit a hyperplane that best separates day 5 than day 6 embryos. All the data was randomly divided into two sets, a training set (70%) and a test set (30%). We performed SVM on the training set, and then we tested the accuracy of this model using the test set.

Last, to understand the conditional dependencies between embryo day, expansion stage and the metabolic parameters, we performed probabilistic graphical models^82,83^ on standardized embryo metabolic parameters 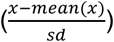, embryo day and stage. A directed arrow was drawn from variable a (embryo day) to variable b (embryo stage) if the probability distribution of a depends on b, and can be written as P(a, b) = P(a)P(b|a). These conditional dependencies can be represented by a DAG. We then computed the percentage of variance explained by both embryo day and stage for each level 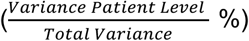, which can represent the effect size of the model^64^.

## Data availability

The authors claim that all relevant data of the findings in this work are provided within the paper and Supplementary Information files. Raw data are provided in the Source Data File.

## Acknowledgments

We would like to acknowledge Becker and Hickl GmbH and Boston Electronics for the loaning of electronic equipment for this research. We also acknowledge the Boston IVF embryologists and clinicians for their assistance during the project, in particular Dr. Emily Seidler and Dr. Alan Penzias and the Needleman lab, at Harvard University especially Brian Leahy, for useful advice.

## Author Contributions

M.V. contributed to the conceptualization of the idea of the experiment, contributed to data acquisition, analysis, and interpretation, and drafted the manuscript. J.S. contributed to data acquisition and critical revision of the manuscript. X.Y. contributed to data analysis and interpretation of the results and critical revision of the manuscript. T.S. contributed to the conceptualization of the idea of the experiment and critical revision of the manuscript. W.C. contributed to data analysis and critical revision of the manuscript. D.S. contributed to the conceptualization of the idea of the experiment and interpretation of the results and critical revision of the manuscript. D.J.N. contributed to the conceptualization of the idea of the experiment and interpretation of the results and critical revision of the manuscript.

