## Supplementary Figures for "Metabolic state of human blastocysts measured by fluorescence lifetime imaging microscopy"

**Movie S1: Time-lapse images of NAD(P)H and FAD+ autofluorescence intensity and average photon arrival time imaging reveal metabolic variations during blastocyst development**. FLIM time-lapse imaging of the autofluorescence of NAD(P)H (top left), and of FAD+ (top right, in grey) and SHG spindle imaging (top right, in magenta) of a human blastocyst throughout 36h of incubation. Average photon arrival times of NAD(P)H (bottom left), and of FAD+ (bottom right) are also shown.

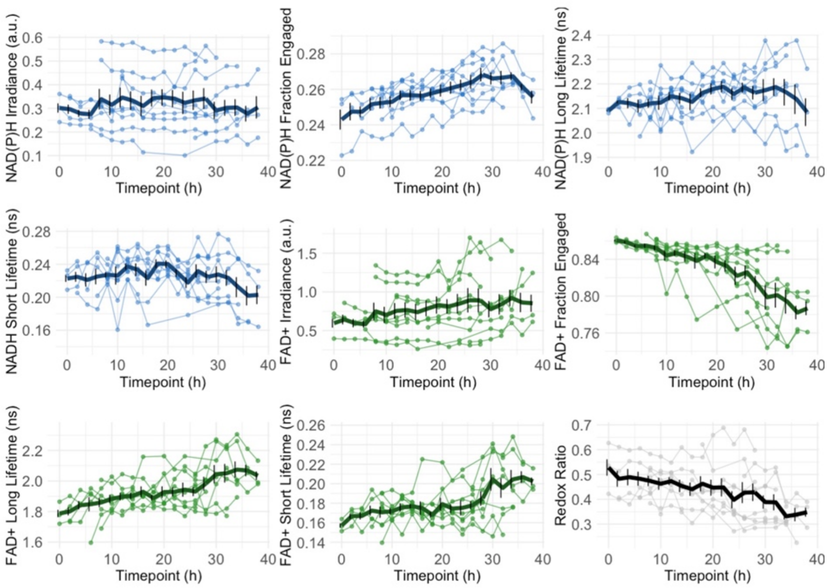


**SI-1: Time curves of metabolic parameters produced through time-lapse FLIM imaging**. These plots display the trajectories of individual embryos (n=10) of NAD(P)H FLIM metabolic parameters (curves in blue), FAD+ metabolic parameters (curves in green) and the redox ratio (in grey). The average trajectories from all the embryos are shown in thick lines with standard error bars. This demonstrates a high reproducibility among all embryos.

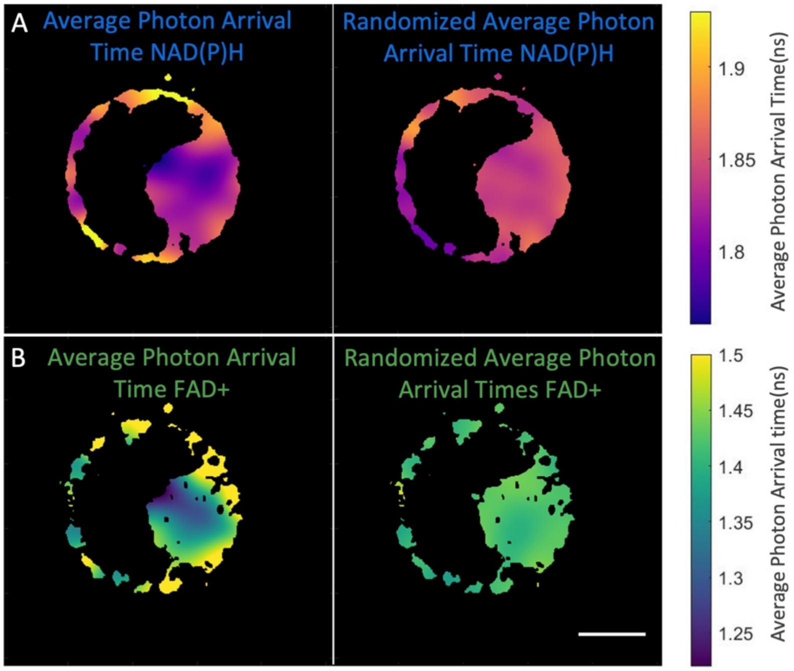


**SI-2: FLIM average photon arrival time images.** (A) Example of an average photon arrival time image of NAD(P)H and (B) FAD+ show the spatial pattern of metabolic signatures between the inner cell mass and trophectoderm (left images). In the right images, the average photon arrival time was homogeneous for randomized photon arrival time images, and do not show a characteristic spatial distribution. Color bars show the average photon arrival time for both NAD(P)H and FAD+ in nanoseconds. Scale bar, 40 μm.

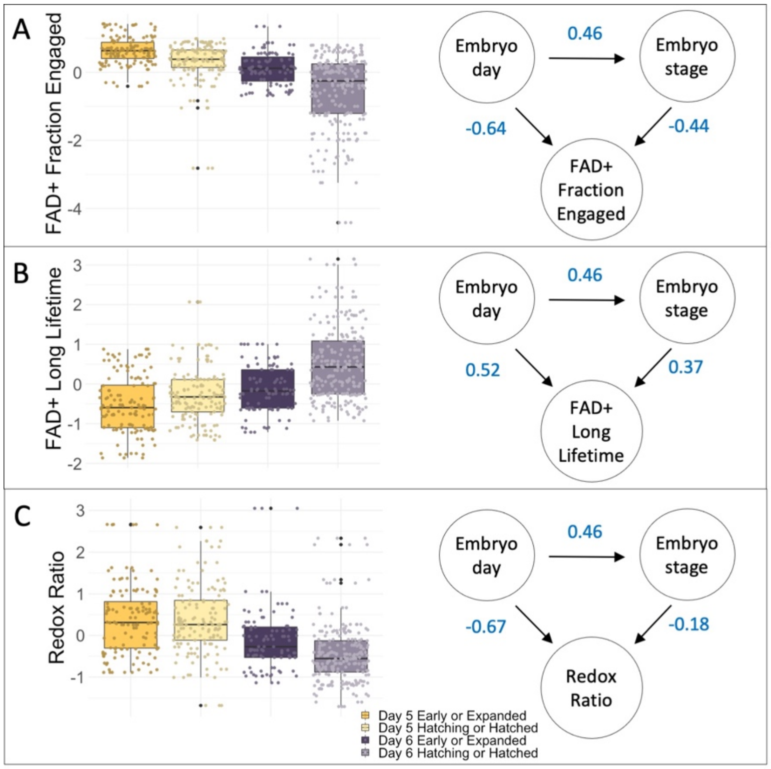


**SI-3. Directed acyclic graphical models of FLIM parameters of human blastocysts and embryo day and expansion stage.** On the left side, the box plots display normalized FAD+ fraction engaged (A), FAD+ long lifetime (B) and redox ratio (C) of 4 categories (Day 5 early/expanded, Day 5 hatching/hatched, Day 6 early/expanded or Day 6 hatching/hatched embryos), showing that both day and stage impact the FLIM parameters. In order to explore the conditional dependencies between embryo day, stage and FLIM parameters we performed probabilistic graphical models. On the right side, DAG of FAD+ fraction engaged (A), FAD+ long lifetime (B) and redox ratio (C) showing that embryo day and stage are correlated and that embryo FLIM parameters are dependent both on embryo day and stage. Numbers in blue represent the $\beta$-coefficient of the multilevel model.
